# A neural circuit mechanism of categorical perception: top-down signaling in the primate cortex

**DOI:** 10.1101/2020.06.15.151506

**Authors:** Bin Min, Daniel P. Bliss, Arup Sarma, David J. Freedman, Xiao-Jing Wang

## Abstract

In contrast to feedforward architecture commonly used in deep networks at the core of today’s AI revolution, the biological cortex is endowed with an abundance of feedback projections. Feedback signaling is often difficult to differentially identify, and its computational roles remain poorly understood. Here, we investigated a cognitive phenomenon, called categorical perception (CP), that reveals the influences of high-level category learning on low-level feature-based perception, as a putative signature of top-down signaling. By examining behavioral data from a visual motion delayed matching experiment in non-human primates, we found that, after categorization training, motion directions closer to (respectively, away from) a category center became more (less) difficult to discriminate. This distance-dependent discrimination performance change along the dimension relevant to the learned categories provides direct evidence for the CP phenomenon. To explain this experimental finding, we developed a neural circuit model that incorporated key neurophysiological findings in visual categorization, working memory and decision making. Our model accounts for the behavioral data indicative of CP, pinpoints its circuit basis, suggests novel experimentally testable predictions and provides a functional explanation for its existence. Our work shows that delayed matching paradigms in non-human primates combined with biologically-based modeling can serve as a promising model system for elucidating the neural mechanisms of CP, as a manifestation of top-down signaling in the cortex.

**Significant Statement:** Categorical perception is a cognitive phenomenon revealing the influences of high-level category learning on low-level feature-based perception. However, its underlying neural mechanisms are largely unknown. Here, we found behavioral evidence for this phenomenon from a visual motion delayed matching experiment in non-human primates. We developed a neural circuit model that can account for this behavioral data, pinpoints its circuit basis, suggests novel experimentally testable predictions and provides a functional explanation for its existence. Our work shows that delayed matching paradigms in non-human primates combined with biologically-based modeling can serve as a promising model system for elucidating the neural mechanisms of categorical perception, as a manifestation of top-down signaling in the cortex.

## Introduction

Categorical perception (CP) is a cognitive phenomenon that reveals complex interplay between analog feature-based perception and discrete categorization (Harnad, 1987). More specifically, category knowledge is thought to warp perception such that differences in appearance between objects that belong to different categories are exaggerated (expansion), while differences within the same category are deemphasized (compression). For example, it was shown that the phonemic categories possessed by adult speakers of English changes these speakers’ perceptual discrimination of consonant-vowel syllables such that intra-category discrimination performance is worse than inter-category discrimination (Liberman et al., 1957). CP has been found in various domains, from phonemic perception in speech (Liberman et al., 1957), to perception of facial expressions (Etcoff and Magee, 1992), to perception of low-level visual features (Goldstone, 1994). Some CP effects arise early in development and persist across adulthood (such as for phonemes in one’s native language), but other CP effects – generally referred to as “learned CP” – can appear after episodes of category learning during adulthood, even for arbitrary, experimenter-defined categories. Given its ubiquity, CP is widely regarded as a key cognitive process that supports the high-level conceptual process through which people organize their world and communicate with each other (Harnad, 1987; Goldstone and Hendrickson, 2010).

Despite the importance of CP to cognition, its underlying neural mechanisms are still largely unknown (Goldstone and Hendrickson, 2010). For instance, it is unclear where in the brain the influence of categorization experience on perceptual processing occurs and how this influence is reflected in both the single neuron and neural population levels. What is lacking in CP studies is a model system that allows researchers to investigate the influences of long-time category learning on perceptual discrimination through simultaneous behavioral assessment and single neuron recordings. This was also indicated in a recent CP study with songbirds (Prather et al., 2009).

Recently, delayed matching experimental paradigms have been established to help unravel the neural mechanisms of visual categorization in non-human primates (Freedman and Assad, 2016). In particular, flexible neuronal categorical representations have been found in parietal and prefrontal areas in monkeys trained to group visual stimuli into arbitrary categories (Freedman and Assad, 2006). This suggests the possibility of using these paradigms to study how the neuronal categorical representations acquired during categorization training influence perceptual processing (i.e., the learned CP effect) in non-human primates. To test this possibility, we examined behavioral data from a visual motion delayed matching experiment that consists of both categorization and discrimination tasks. We found that, during the motion direction discrimination task after categorization training, motion directions closer to the category center became more difficult to discriminate than those farther away from the category center, a signature of CP phenomenon.

To explain this experimental finding, based upon previous biophysical models (Engel and Wang, 2011; Engel et al., 2015), we developed a neural circuit model of CP that leveraged existing key neurophysiological findings in visual categorization (Freedman and Assad, 2016), working memory (Leavitt et al., 2017) and perceptual decision making (Gold and Shadlen, 2007). By hypothesizing that the category knowledge acquired during categorization training entered into the discrimination circuit through feedback projections, this model was shown to be capable of accounting for the behavioral data in our experiment. One key testable prediction from this model is the existence of a mixture area that integrates feedforward sensory input and feedback category input. Furthermore, by showing that category knowledge acquired during categorization training can help improve perceptual stability in noisy environments, our work provided a functional explanation for the existence of CP phenomenon. In summary, by providing behavioral evidence for learned CP and developing a computational framework that both integrates disparate neurophysiological findings and allows us to make specific testable predictions for further experiments, our work suggests that delayed matching paradigms in non-human primates can serve as a promising model system for elucidating the neural mechanisms of learned CP phenomenon.

## Materials and Methods

### Behavioral tasks and stimulus display

Monkeys performed delayed match-to-sample (DMS) and delayed match-to-category (DMC) tasks using 360° of motion directions as sample and test stimuli. Monkeys released a manual lever to indicate whether a test stimulus was the same direction (DMS) or same category (DMC) as a previously presented sample. Monkeys were required to maintain fixation within 2.0-2.5° of a 0.2° fixation point throughout the entire trial. Sample and test stimuli were 9.0° diameter circular patches of high-contrast, 100% coherent random dots. Dots moved at 12° per second and were displayed at a frame rate of 75 Hz. During both the DMS and DMC tasks, twelve evenly spaced sample directions were used as sample stimuli. During the DMS task, sample and non-matching test stimuli were separated by either 45°, 60° or 75° to keep monkeys’ performance on the task above chance but below complete certainty. During the DMC task, test stimuli were chosen randomly from the same directions as the sample stimuli. In task switching, a trial also began with the onset of a fixation spot. Monkeys were required to maintain gaze within 2.0° of a fixation point throughout the trial. Depending on the fixation spot color (magenta or green), monkeys were instructed to perform different tasks (DMC or DMS, respectively). Stimulus presentation, task events, rewards, and behavioral data acquisition were accomplished using an Intel-based PC equipped with MonkeyLogic software running in MATLAB. Gaze positions were measured and recorded at a sampling rate of 1.0 kHz using an EyeLink 1000 optical eye tracker (SR Research). Visual stimuli were presented on a 21” color CRT monitor (1,280 × 1,024 resolution, 75 Hz refresh rate, 47-cm viewing distance).

### Behavioral training

Monkeys D and H (male *Macaca mulatta*, 9.0-12.0 kg, 7-9 years old) were each trained on the motion DMS task for approximately 1 year (about 1,400 correct trials/session). Monkeys were trained each day on 12 unique sample directions, and test directions were chosen to include a wide range of angular differences between non-matching sample and test directions. Early DMS training sessions focused on large angular differences between sample and test directions (for example, 90–180°) until monkeys’ accuracy reached a high level. Middle to late stage training sessions focused on more challenging sample-test differences (for example, 5–30°), and DMS training continued until the monkeys’ performance improved and stabilized. DMC task training of Monkeys D and H was conducted in two stages. The first was the mid-training DMC stage (104 and 70 sessions in monkeys D and H, respectively) in which the category boundary was introduced and the monkeys were rewarded for indicating (with a lever release) whether the test stimulus was in the same category as the sample. In this training stage, the 12 sample directions used for training were shown with equal frequency during each session, and a wide range of sample-test differences were used. In the second DMC training stage (78 and 65 sessions in monkeys D and H), we over-emphasized near-boundary (that is, 15° from boundary) sample stimuli for which the monkeys had shown lower performance in the middle training stage. This training stage continued until the monkeys’ performance for near-boundary stimuli improved and stabilized. Monkeys D and H were each trained on task switching more than 6 months. After that, the monkeys performed both tasks with high accuracy as shown in Fig.1.

**Figure 1:**
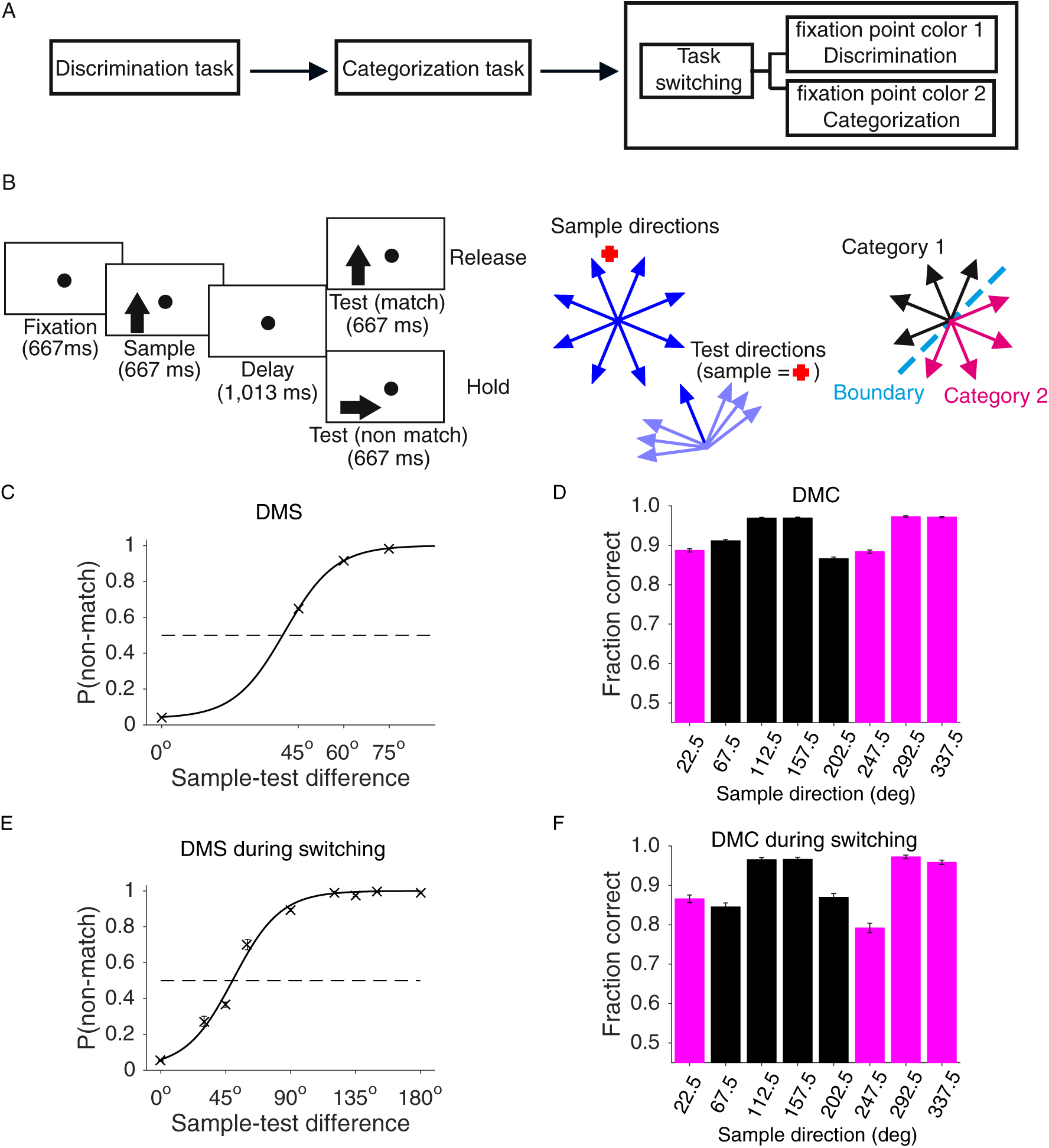
A visual motion delayed matching experiment in non-human primates. **(A-B)** Monkeys were trained to perform a discrimination task, then a categorization task, finally task switching between the two. Both discrimination and categorization tasks are of the delayed matching type, in which sample stimulus, consisting of moving random dots, is followed by a delay and then a test stimulus, and monkeys were required to report (by releasing a lever) if the test stimulus was a match to the previous sample stimulus. In the discrimination task, termed delayed match-to-category (DMS) task below, match means the motion directions of sample and test stimuli are exactly same, while in the categorization task, termed as delayed match-to-category (DMC) task below, match means sample and test stimuli belong to the same predefined category. In task switching, monkeys were required to perform either the DMS or DMC tasks depending on the fixation point color. (**C, E**) The DMS task performance before (**C**) and after (**E**) DMC training. (**D, F**) The DMC task performance before (**D**) and during (**F**) task switching. Error bars indicate s.e.m.

All surgical and experimental procedures followed the University of Chicago’s Animal Care and Use Committee and US National Institutes of Health guidelines. Monkeys were housed in individual cages under a 12-h light/dark cycle. Behavioral training was conducted during the light portion of the cycle.

### Task protocol in the simulation

In our simulation, an ABA task scheme (Fig. 3A) was used, in which A is a DMS task and B is a categorization task. The DMS task is the same as the one used in the experiment. In the categorization task here, one randomly chosen stimulus was presented, and a neural network was trained to learn its category membership through trial and error. This is different from the DMC task in the experiment, in which there were two stimuli to be compared with each other.

**Figure 2:**
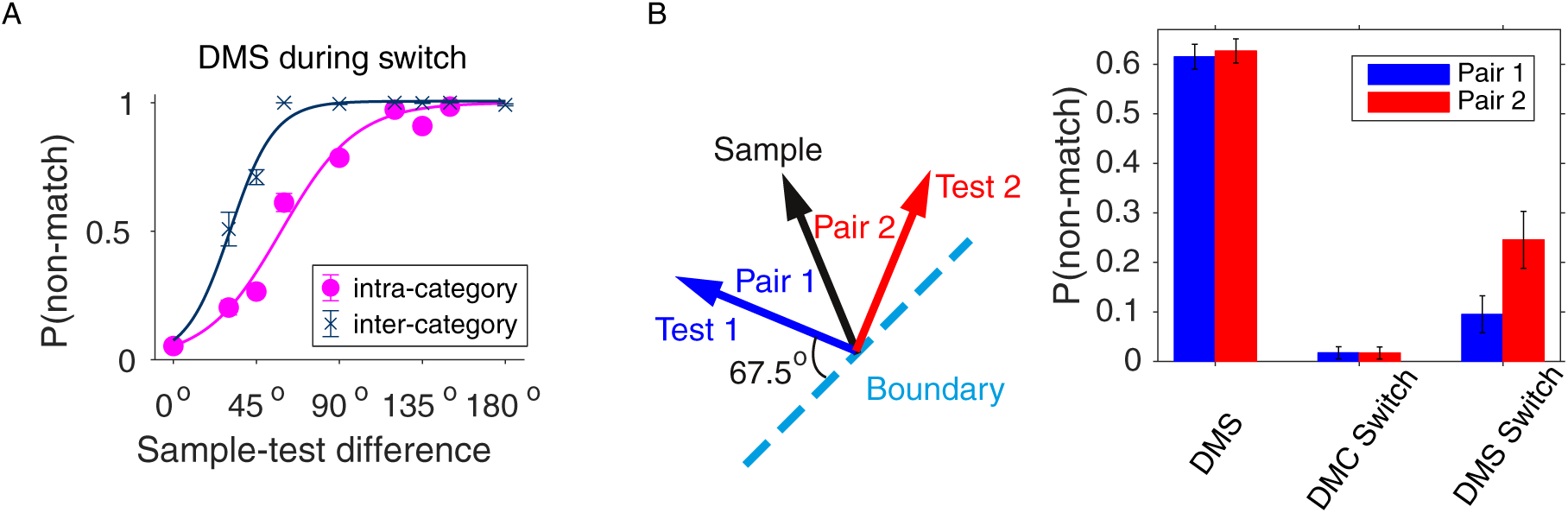
Behavioral evidence for the CP effect in this experiment. **(A)** Behavioral performance for intra- and inter-category groups as a function of sample-test directional difference. The intra-category pairs are those with sample and test stimuli in the same category, while the inter-category pairs are those with sample and test stimuli belonging to different categories. (**B**) Comparison between two intra-category sample-test pairs. These two pairs share the same sample direction and have the same sample-test directional difference (45°) (left panel). Before DMC training, there was no significant DMS performance difference between these two pairs, *p* = 0.369 (two-proportion z-test). During task switching, there also was no significant DMC performance difference between these two pairs, *p* = 0.507 (two-proportion z-test). However, there was a significant DMS performance difference between these two pairs during task switching, *p* = 0.014 (two-proportion z-test). Error bars indicate s.e.m.

**Figure 3:**
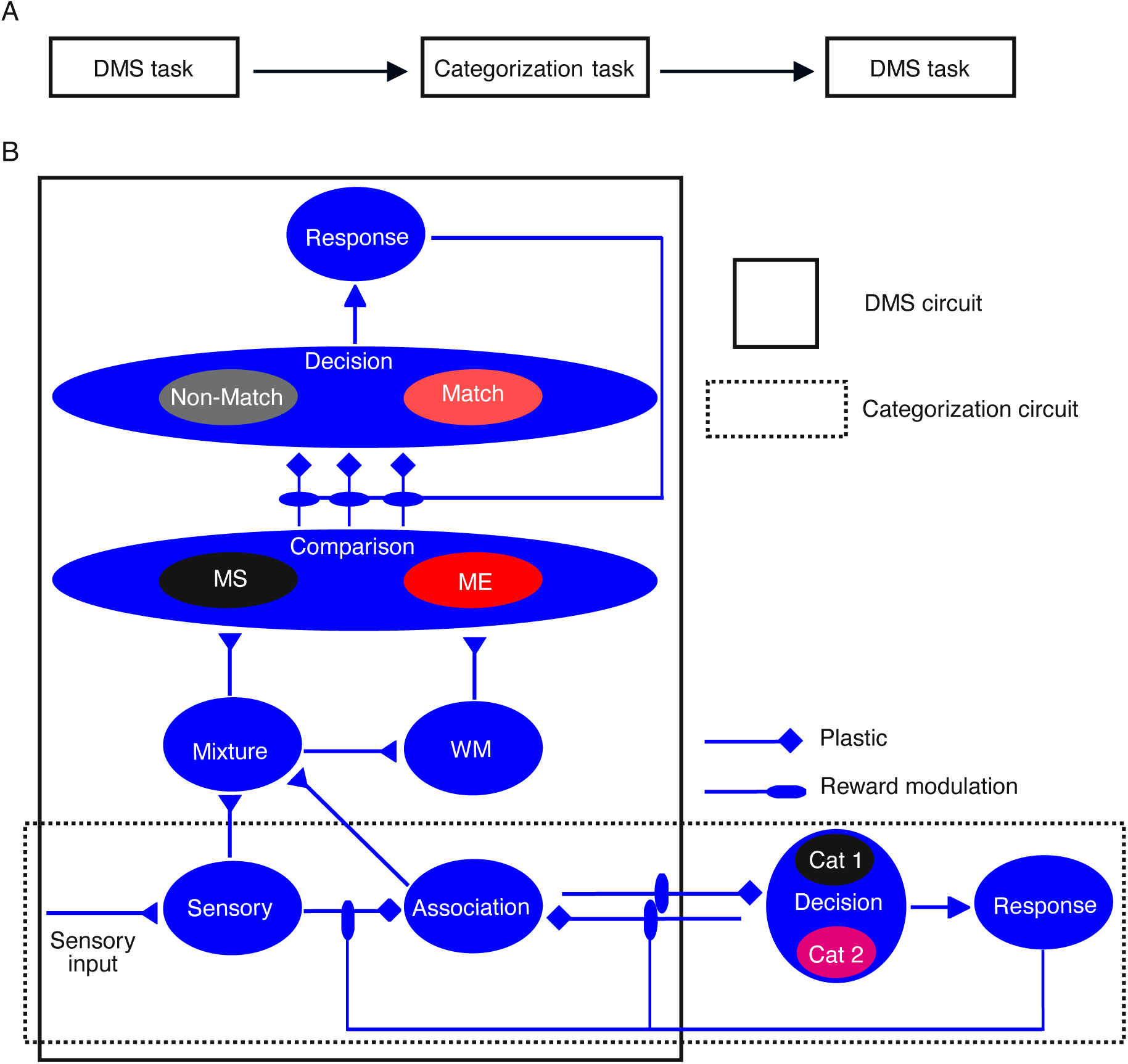
The architecture of our neural circuit model. **(A)** Schematic of the task structure we used in the neural circuit simulation. Our neural circuit is trained to perform an ABA task scheme in which A is a DMS task and B is a categorization task. The DMS task here is the same as the one used in the experiment. In the categorization task, one randomly chosen stimulus is presented, and the neural circuit is trained to learn its category membership through trial and error. (**B**) Organization diagram of the circuit model. Solid black box: the DMS circuit; dashed black box: the categorization circuit. The two are linked primarily through the projection from the association area to the mixture area. The DMS circuit consists of four functional modules, including sensory representation (sensory, association and mixture), persistent working memory (WM), comparison (MS and ME) and decision (non-match and match) modules. The synaptic strengths of connections from the comparison module to the decision module undergo reward-modulated Hebbian plasticity at the end of each DMS trial. The categorization circuit consists of sensory, association and decision areas. The synaptic strengths of all inter-area (sensory-to-association, association-to-decision, association-to-decision) connections undergo reward-modulated Hebbian plasticity at the end of each categorization trial.

### Description of the computational framework

The full computational model consists of a categorization circuit and a DMS circuit. In the categorization circuit, there are three functional areas, including sensory (S), association (A) and decision (D, including population C1 and C2), of which the sensory and association areas are shared with the DMS circuit. In the DMS circuit, there are four modules for sensory representation (including sensory, association and mixture areas), working memory (WM), comparison (including population match-enhancement [ME] and match-suppression [MS]) and decision (including population match [M] and non-match [NM]).

#### The categorization circuit

This categorization circuit is the same as the one developed in the previous work (Engel et al., 2015). All areas, including sensory, association and decision, are strongly recurrent networks with dynamics governed by local excitation and feedback inhibition. In simulations, we used a reduced mean-field model that has been shown to reproduce neural activity of a full spiking neural network (Wong and Wang, 2006). The dynamics of each excitatory population in the mean-field model is described by a single variable *s* representing the fraction of activated N-methyl-D-aspartate receptor conductance, governed by

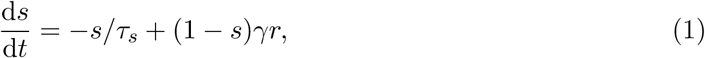

where *γ* = 0.641 and *τ*_*s*_ = 60 ms. The firing rate *r* is a function of the total synaptic current *I*:

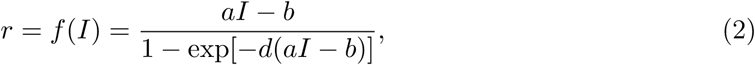

with *a* = 270 Hz nA^−1^, *b* = 108 Hz and *d* = 0.154 s.

The total synaptic current *I* consists of recurrent and noisy components, *I* = *I*_*r*_ + *I*_*n*_. Recurrent input to a neuron *i* in the population A originating from the population B (population B can be either the same as or different from population A) reads:

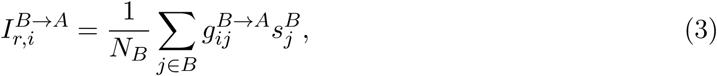

where 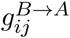 is the synaptic coupling between the neuron *j* in the population B and the neuron *i* in the population A. The current is normalized by the number of presynaptic neurons *N*_*B*_. Noisy current replicates background synaptic input and obeys:

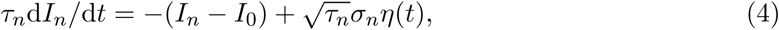

where η(*t*) is a white Gaussian noise, *τ*_*n*_ = 2 ms, σ_*n*_ = 0.009 nA, 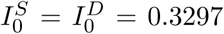 nA and 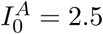nA.

The sensory and association areas were each simulated by 128 discrete units with equally spaced preferred directions from 0° to 360°. Within each area, the synaptic couplings *g*_*ij*_ between neurons with preferred directions *θ*_*i*_ and *θ*_*j*_ have a periodic Gaussian profile:

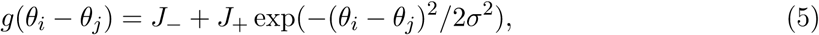

where *σ* = 43.2°. Parameters *J*_−_ and *J*_+_ determine the amount of recurrent excitation and inhibition. In sensory and association areas, 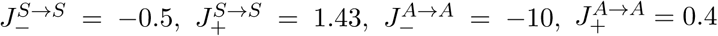 nA.

The decision area consists of two populations (C1 and C2) respectively selective for category 1 (Cat 1) and category 2 (Cat 2), which are driven by the association neurons. When stimulated, activities of the C1 and C2 populations diverge according to winner-take-all dynamics. This behavior is attained through global inhibition and structured recurrent excitation within the decision area: 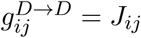 with 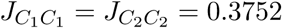 nA and 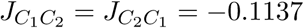 nA.

During the categorization task, all synapses connecting three areas (from sensory to association, and between association and decision neurons) are plastic and excitatory. Synaptic strengths of plastic connections are expressed as *g*_*ij*_ = *g*_*max*_*c*_*ij*_, where *g*_*max*_ is the maximal connection strength and *c*_*ij*_ is bounded between 0 and 1, and represents the fraction of potentiated synapses between neurons *i* and *j*. At the end of each trial, all *c*_*ij*_ are updated according to the Hebbian plasticity rule modulated by the reward prediction error as specified below:

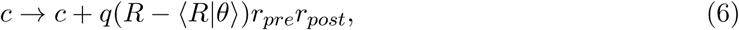

where *r*_*pre*_ and *r*_*post*_ are the trial-average firing rates of pre- and post-synaptic neurons, *q* is the learning rate parameter, *R* is the reward received on each trial (1 or 0 for correct and incorrect decisions, respectively), *θ* stands for a motion direction stimulus and ⟨ *R*|*θ*⟩ is a stimulus-specific reward expectation.

Plastic synapses between sensory and association neurons 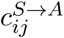 are initialized with the periodic Gaussian profile as in equation (5) with 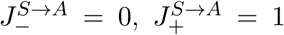. Plastic synapses between association and decision neurons (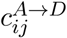 and 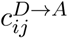) are initialized randomly from a uniform distribution on [0.25, 0.75]. The maximal connection strengths of plastic synapses are 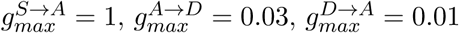 nA.

Each simulation trial started with a 200-ms pre-stimulus period (no external input), followed by a 1-s presentation of a motion direction stimulus and then by a 500-ms inter-trial interval. When a motion direction stimulus *θ*_*s*_ is presented, neurons in the sensory population receive additional input current *I*_*s*_ that depends on the neuron’s preferred direction *θ*:

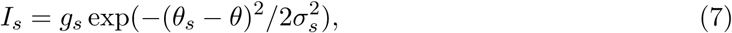

where *σ*_*s*_ = 43.2°and *g*_*s*_ = 0.1 nA. Neurons in the decision circuit receive a non-selective gating current of 0.01 nA during the stimulus period, which sets the circuit in the decision-making regime, and a brief −0.08 nA reset current during the first 300 ms of the inter-trial interval, which represents the corollary discharge and resets activity to the spontaneous level.

The model’s response on each trial is determined by comparing firing rates of two decision populations with a 20-Hz threshold during the last 25-ms of the stimulus period. The response is considered invalid if both or neither population reach threshold, or either population reaches threshold before the stimulus onset. Across trials, choices of the decision area are stochastic and are characterized by a sigmoidal dependence of the probability of choice *C*_1_ on the difference Δ*I* in synaptic input currents to two competing populations. Reward equals *R* = 1 on valid correct trials, *R* = 0 on valid incorrect trials and no plasticity is triggered on invalid trials.

#### The DMS circuit

The DMS circuit is built upon the one developed in the previous work (Engel and Wang, 2011). This circuit shares the sensory and association areas with the categorization circuit. We describe the remaining populations in the DMS circuit here, including the mixture, WM, ME, MS, M and NM populations. All of these are strongly recurrent networks with dynamics governed by local excitation and feedback inhibition. More specifically, the dynamics of each population is described by the mean-field equation (1) with F-I curve (2). The total synaptic current *I* in equation (2) consists of recurrent, external sensory and noisy components, *I* = *I*_*r*_ + *I*_*s*_ + *I*_*n*_. Recurrent input to a neuron *i* in the population A originating from the population B is specified by equation (3).

The mixture area and the WM module were each simulated by 128 discrete units with equally spaced preferred directions from 0° to 360°. Within each area/module, the synaptic couplings *g*_*ij*_ between neurons with preferred directions *θ*_*i*_ and *θ*_*j*_ follow the same profile specified by equation (5) with *σ* = 43.2°. In the mixture area and the WM module, 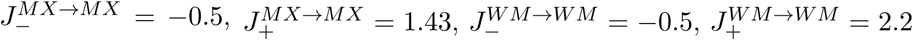 nA. The stronger recurrent excitation enables persistent firing within the WM module during the delay period. In addition, the mixture area received inputs from both the sensory and association areas. The synaptic couplings between neurons in the mixture area with preferred directions *θ*_*i*_ and neurons in both the sensory and association areas with preferred directions *θ*_*j*_ follow the same profile specified by equation (5) with 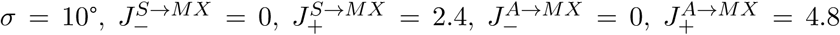 nA. The WM module receives input from the mixture area with the Gaussian profile specified by equation (5) with 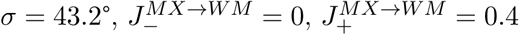 nA.

The ME and MS populations were each simulated by 128 discrete units with equally spaced preferred directions from 0° to 360°. ME neurons receive excitation from the WM module with the Gaussian profile specified by equation (5) with 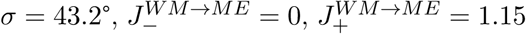 nA. In contrast, MS neurons do not receive any top–down input: 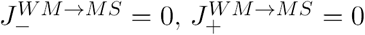 nA. We assume that excitatory conductances of the ME cells are weakened by a factor *α* = 0.975 because of a homeostatic mechanism acting to compensate for the top-down excitation in these cells. This homeostatic mechanism is operating on a very slow timescale, so that the value of *α* = 0.975 is held constant in all simulations. Within the comparison module that consists of the ME and MS populations, the synaptic couplings *g*_*ij*_ between neurons with preferred directions *θ*_*i*_ and *θ*_*j*_ follow the same profile specified by equation (5) with *σ* = 43.2°, *J*_−_ = −8.5 nA, 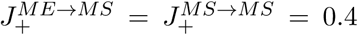 nA and 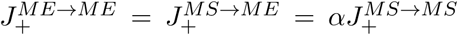. Both the ME and MS neurons receive excitation from the mixture area with the Gaussian profile specified by equation (5) with 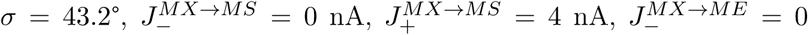 nA, 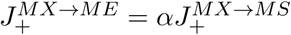.

We assume that sensory signals reach the WM module only when attention is directed to store the sample in the WM. Signals from the test stimuli do not reach the WM circuit. In all simulations, sensory stimuli were presented for 0.6 s using the same Gaussian current injection as in the category task. Sample and test stimuli were separated by a 1-s delay.

Noisy background synaptic input obeys equation (4) with *τ*_*n*_ = 2 ms, *σ*_*n*_ = 0.009 nA, 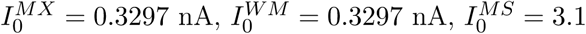 nA, and 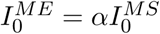.

The ME and MS neurons have an additional current *I*_*a*_ mimicking the spike rate adaptation as follows: *I*^{*ME,MS*}^ = *I*_*r*_ + *I*_*s*_ + *I*_*n*_ + *I*_*a*_, whereby *I*_*a*_ = *g*_*a*_*s*_*a*_ and *g*_*a*_ = 0.003 nA. The dynamics of *s*_*a*_ follows *ds*_*a*_*/dt* = −*s*_*a*_*/τ*_*a*_ + *r*, with *τ*_*a*_ = 10 s.

The activities of the ME and MS neurons are pooled by the decision circuit with two competing neural populations selective for choice “match” and “non-match” (See Fig. 3B). When stimulated, activities of the two populations diverge according to winner-take-all dynamics, and the decision of the model is determined by the population with a higher activity. Across trials, the stochastic choice behavior of the decision circuit is characterized by a sigmoidal dependence of the probability P to choose non-match on the difference Δ*I* in synaptic input currents to the non-match and match pool. More specifically, given synaptic input currents to the non-match and match pools from the ME and MS populations,

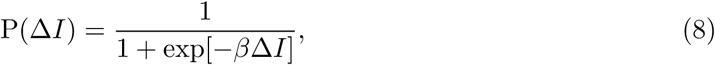

where P(Δ*I*) is the fraction of choosing non-match and Δ*I* is the difference of synaptic input currents to the non-match and match pools (Soltani and Wang, 2006). We used β = 200 nA^−1^.

The synapses connecting comparison neurons with the decision neurons are plastic. Each pair of presynaptic and postsynaptic cells is connected by a set of binary synapses that are in either a potentiated or a depressed state. The fraction 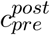 of synapses in the potentiated state quantifies the strength of synaptic connection. Input currents to the match and non-match populations are expressed through the synaptic strengths as 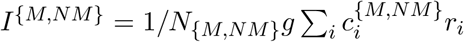, where the sum goes through all neurons in the comparison network, *r*_*i*_ are their firing rates, and *g* = 2 nA/Hz. At the end of each trial, all synapses onto the chosen population (match or non-match) are updated according to a reward-dependent Hebbian plasticity rule. If the choice of the model is rewarded, the synapses are potentiated (i.e., the synapses in the depressed state make a transition to the potentiated state with the rate *q*_0_ · *q*(*r*) referred to as the learning rate) as follows:

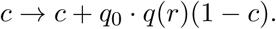

If the choice of the model is not rewarded, the synapses are depressed as follows:

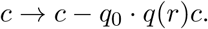

The maximal learning rate *q*_0_ determines the speed of learning. The learning rate depends on the presynaptic firing rate:

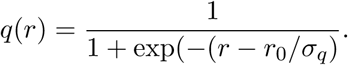

We used *r*_0_ = 8 Hz, *σ* = 4 Hz and *q*_0_ = 10^−3^.

#### Category-tuning index (CTI)

The CTI of each neuron is defined as the difference between the between-category and within-category differences divided by their sum (Freedman and Assad, 2006), in which the between-category (within-category) difference is the difference in firing rate between pairs of directions in different categories (the same category). Values of CTI can vary from 1 (larger differences in neural activity between pairs of directions in different categories) to −1 (larger differences in neural activity between pairs of directions in the same category). A CTI value of 0 indicates the same difference in firing rate between and within categories.

## Results

### Behavioral evidence for CP in non-human primates

In our experiment, monkeys were trained to perform a discrimination task first, then a categorization task, and finally task switching where they switched between discrimination and categorization depending on a task cue (fixation point color) (Fig. 1A). Both the discrimination and categorization tasks were of the delayed matching type, in which a sample stimulus, consisting of moving random dots, was presented first and followed by a delay and then a test stimulus, and monkeys were required to report if the test stimulus was a “match” to the previous sample stimulus by releasing a manual touch bar (Fig. 1B). During the discrimination task, match meant that the motion directions of sample and test stimuli were exactly same, while during the categorization task, match meant that sample and test stimuli belonged to the same category whose boundary was arbitrarily defined by the experimenters (and not presented explicitly to the subjects). The discrimination and categorization tasks here will be termed as delayed match-to-sample (DMS) and delayed match-to-category (DMC) tasks, respectively.

Monkeys were extensively trained during DMS, DMC and task switching. During the DMS task, behavioral performance was greater than chance (50%) when sample and test stimuli were 45° apart, and greater than 85% correct on all of the other non-match and match conditions (Fig. 1C). During the DMC task, behavioral performance was *>* 85% correct for the four sample directions that were 22.5° from the category boundary and *>* 90% correct for the four directions that were 67.5° from the boundary (Fig. 1D). During task switching, monkeys performed well, with the DMS performance greater than 85% correct when sample and test stimuli were 90° apart (Fig. 1E), and the DMC performance *>* 75% correct for the four sample directions that were 22.5° from the category boundary and *>* 90% correct for the four directions that were 67.5° from the boundary (Fig. 1F).

Although monkeys generally performed well during task switching, DMS performance during task switching was impaired relative to performance before DMC training. For example, DMS performance during task switching was about 40% when sample and test stimuli were 45° apart (Fig. 1E), significantly worse than that (above 60%) during the DMS task before DMC training (Fig. 1C), *p <* 10^−68^ (two-proportion z-test). Some impairment was expected due to the higher cognitive demands associated with task switching. To address this possibility, we divided sample-test pairs into two groups – one intra-category group with sample and test stimuli in the same category and one inter-category group with sample and test stimuli belonging to different categories. We found that DMS performance for the intra-category pairs during task switching was much worse than that for the inter-category pairs (Fig. 2A). This difference cannot be explained by the higher cognitive demands during task switching, as these demands should be constant regardless of the category membership of stimulus pairs. Hence, the difference in performance between intra-category and inter-category stimulus pairs indicated the existence of complex interaction between categorization and discrimination.

One possibility of this interaction could be that during task switching there was a fraction of DMS trials in which monkeys erroneously applied the DMC match rule. For each sample-test pair, we defined a probability of rule-confusion as the proportion of DMS trials in which monkeys erroneously applied the DMC match rule. Given the same sample-test difference value, following the DMC match rule will lead to higher probability of choosing match for the intra-category pairs than the inter-category pairs. This may explain the DMS performance discrepancy between the intra- and inter-category groups during task switching (Fig. 2A). However, such rule confusion would not qualify as genuine CP, conventionally defined. To assess this possibility, we compared two intra-category sample-test pairs which have the following properties: (1) these two pairs share the same sample stimulus, (2) sample and test stimuli were 45° apart for both pairs, (3) before DMC training, there was no significant DMS performance difference between these two pairs, *p* = 0.369 (two-proportion z-test), (4) during task switching, there was no significant DMC performance difference between these two pairs, *p* = 0.507 (two-proportion z-test) (Fig. 2B). In this experiment, as the rule cue was explicitly presented during the whole trial period, given the same sample stimulus, the probability of rule-confusion, if present, should be independent of the incoming test stimuli, therefore the same for these two pairs. Then, it was expected that during task switching, there also would be no significant DMS performance difference between these two pairs. However, we found that during task switching, the DMS performance of the pair closer to the category center (pair 1) actually was significantly worse than that of the pair farther from the category center (pair 2), *p* = 0.014 (two-proportion z-test) (Fig. 2B). Following this logic, the rule-confusion hypothesis was disconfirmed, lending support that pair 1 becomes harder to discriminate than pair 2 as a result of categorization training.

Taken together, our data analysis suggests that in DMS trials during task switching motion directions closer to the category center became more difficult to discriminate than those farther away from the category center, demonstrating the CP effect in this experiment.

### Architecture of neural circuit model

To gain insights into neural mechanisms underlying the CP effect in this experiment, we developed a neural circuit model. The major issue we addressed in this neural circuit model is where the influence of categorization experience on perceptual processing occurs and how this influence is reflected both in single neurons and at the neural population level. To this end, an “ABA” task scheme (Fig. 3A), in which “A” is the DMS task and “B” is a simplified categorization task, was used in our model simulation. In this simplified “one-interval” categorization task, instead of presentation of two stimuli to be compared with each other in each trial, as in the DMC task in the experiment, there was only one stimulus to be categorized. Another noticeable difference from the experiment is that the same DMS task A, instead of task switching, was used after category learning (task B).

Our network model is built upon two previous models, one for DMS (task A) (Engel and Wang, 2011), the other for categorization (task B) (Engel et al., 2015) (for details, see **Materials and Methods**). Similar to previous biophysical models (Engel et al., 2015; Compte et al., 2000; Wang, 2002; Wong and Wang, 2006; Engel and Wang, 2011), both of the categorization and DMS circuits comprise several functional areas/modules (Fig. 3B). The categorization circuit consists of sensory, association and decision areas, each of which is associated with a functional role in the categorization task and importantly is constrained by physiological findings. Similarly, in the DMS circuit, there are four modules for sensory representation (including sensory, association and mixture areas), working memory (WM), comparison and decision. Each module is associated with a functional role in the DMS task and importantly is also constrained by physiological findings. Our circuit model can be regarded as a minimal biophysical model. We did not intend to explain every aspect of experimental results but aimed at accounting for salient experimental findings. The two circuits are linked by a connection from the association area in the categorization circuit to the mixture area in the DMS circuit.

More specifically, in the categorization circuit, the sensory area – which corresponds to the middle temporal (MT) area in cortex, for the case of a motion direction stimulus – encodes stimuli with a bell-shaped activation profile, arising from bottom-up sensory input and structured recurrent excitation (Engel and Wang, 2011) (Fig. 4). Initially, the association area (lateral intraparietal [LIP] area) inherits direction selectivity from upstream motion processing (Fig. 4), which was achieved by setting stronger synaptic weights between sensory and association neurons with more similar preferred directions. As was shown in previous work (Engel et al., 2015), reward-modulated Hebbian plasticity for synaptic connections from sensory neurons to association neurons caused the categorization circuit to develop stable category representation in this association area over the course of categorization training in our simulations (Fig. 4). Importantly, the emergence of this neuronal category representation in the association area recapitulated the presence of category-tuned neurons in the LIP area during a visual motion categorization experiment (Freedman and Assad, 2006). The decision area comprises two competing decision pools, firing at higher rates for the two respective category decisions (Wong and Wang, 2006).

**Figure 4:**
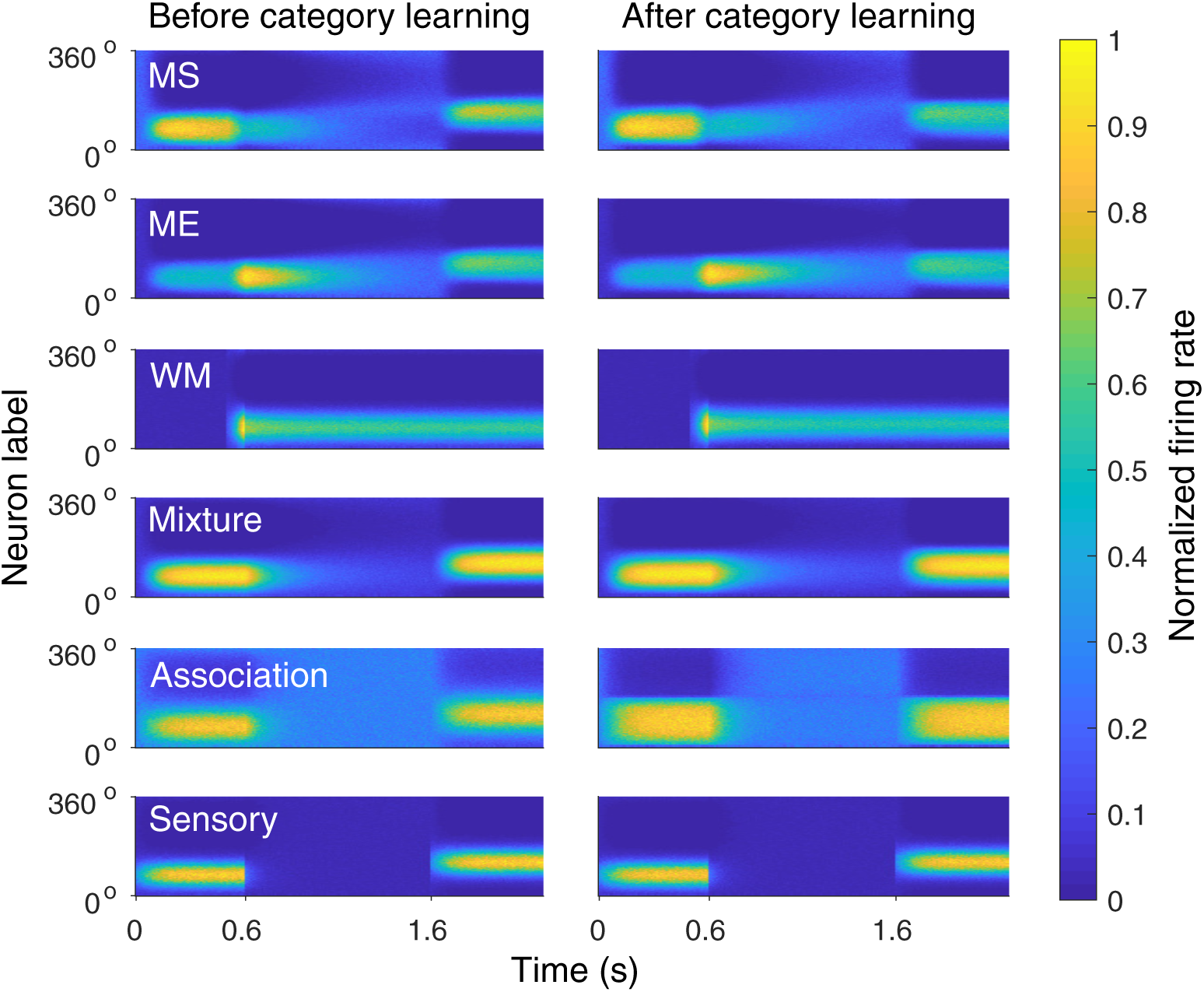
An example neural activity in the model during DMS task. The left and right panels correspond to the cases before and after categorization training, respectively. A sample stimulus (67.5°) was presented for 0.6 s. After 1 s delay, a test stimulus (112.5°) was presented for another 0.6 s. Before category learning (with the category boundary at 0° and 180°), the sensory, association and mixture areas showed direction-tuned responses in their spatiotemporal activity patterns during both sample and test periods. After category learning, the association area showed a category-tuned response in its spatiotemporal activity pattern during both sample and test periods. x axis, time from sample onset; y axis, neurons arranged and labelled by their preferred directions before category learning; firing rates normalized by the maximum firing rate in each area/module are color-coded.

In the DMS circuit, in the sensory representation module, there is an additional mixture area that integrates bottom-up input from the sensory area and top-down input from the association area. Although the mixture area is not strictly needed for the model to perform the DMS task, its inclusion matches the organization of primate cortex, where medial superior temporal (MST) area is positioned between MT and LIP and is strongly connected to both. Synaptic weights were set to be stronger between sensory and mixture neurons with more similar initial preferred directions. The same connectivity pattern was also used for synaptic weights between association and mixture neurons. As a result, during our simulations, the mixture area was capable of encoding the motion direction of incoming stimuli before categorization training (Fig. 4). As neuronal category representation emerged in the association area over the course of categorization training, the top-down input from the association area enabled the mixture area to encode category information. In particular, under this setting, we were able to control the strength of neuronal category selectivity in the mixture area by tuning the ratio of the association-to-mixture synaptic weight to the sensory-to-mixture synaptic weight.

In the downstream WM module, neurons receive input from the mixture area at the end of the sample period. The major difference between this WM module and areas in the sensory representation module is the strength of recurrent excitation. While sample information decayed away over tens of milli-seconds after sample stimulus removal in areas in the sensory representation module, the stronger recurrent excitation enabled this WM module to store sample information through synaptic reverberation during the delay period that lasted one second in the DMS task here (Compte et al., 2000). The comparison module consisted of two functional populations – one match-enhancement (ME) population and one match-suppression (MS) population. Neurons in the ME population showed higher firing response for match sample-test pairs than non-match ones, while neurons in the MS population showed higher firing response for nonmatch pairs. In our model, this was achieved through inhibition-dominated recurrent dynamics and heterogeneous top-down excitation from the WM module (Engel and Wang, 2011). In the last neural processing stage of this DMS circuit, a decision module, comprising two competing decision pools – one match pool and one non-match pool, was used (Wang, 2002; Wong and Wang, 2006).

### Neural representations of the sample stimulus during the DMS task in the model

To study functional roles of different areas/modules in the DMS circuit model (Fig. 5A), we visualized the neuronal representations of sample stimuli with firing rate heat maps (Fig. 5B). During the DMS task after categorization training, as the sample direction varies, the association area showed a nearly binary change in the neural firing pattern when the category boundary was crossed, while the sensory area exhibited a gradual change across stimulus directions. Thus, category representation emerged in the association area after categorization training. The mixture area showed a firing pattern intermediate between a gradual change within each category and an abrupt change across the category boundary. This is because the mixture area integrated inputs from the association and sensory areas. As information in the WM module was inherited from the mixture area, the WM module also showed a similar mixture of category and feature signals.

**Figure 5:**
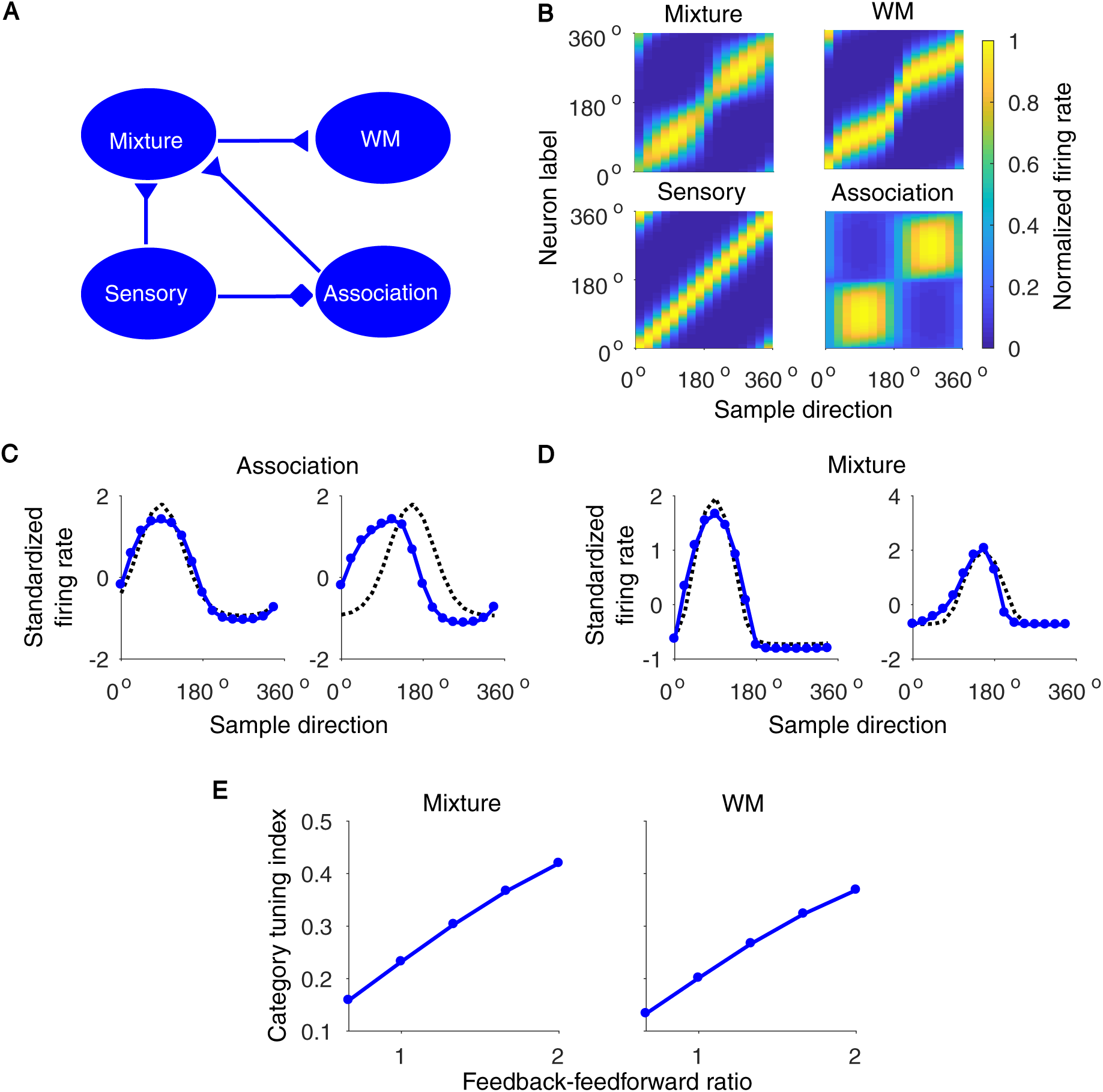
Neural representations of sample stimulus in the model during DMS task after category learning. **(A)** Schematics of the connections between different areas/modules. (**B**) Tuning profiles of different areas/modules. x-axis, stimulus motion direction; y-axis, neurons arranged and labelled by their preferred directions before category learning; firing rates that are normalized by the maximum firing rate of each module are color-coded. (**C-D**) Tuning profiles of two example neurons before (black dashed line) and after (blue solid line) category learning in the association and mixture areas. (**E**) The average category-tuning index of neurons in the mixture area and the WM module as a function of the feedback-feedforward ratio. Here, feedback-feedforward ratio is defined as the ratio of the association-to-mixture synaptic weight to the average sensory-to-mixture synaptic weight.

To better quantify changes in the responses of single neurons after categorization training, we examined individual neuron tuning curves. We found that neurons in the association area showed two interesting trends, consistent with previous findings (Engel et al., 2015). One was tuning curve broadening that occurred for neurons with initial preferred directions close to the category center (Fig. 5C, left panel). The other was tuning curve shifting that occurred for neurons with initial preferred directions close to the category boundary (Fig. 5C, right panel). These two trends also existed in the mixture area but to a lesser degree (Fig. 5D). To quantify how much category information was encoded in the mixture area and the WM module, we computed the average category-tuning index (CTI) (Freedman and Assad, 2006). The average CTI of each population measured how strong the population is tuned to different categories (for details, see Materials and Methods). We also defined a feedback-feedforward ratio as the ratio of the association-to-mixture feedback projection strength to the sensory-to-mixture feedforward projection strength. We plotted the CTI of neurons in different areas/modules as a function of the feedback-feedforward ratio. When the feedback-feedforward ratio is low, categorical information sent to the mixture area is insignificant compared to the feedforward motion direction information this area receives. Increasing this ratio would be predicted to increase the CTI. As expected, the average CTI for both the mixture area and the WM module increases as the feedback-feedforward ratio increases (Fig. 5E). Therefore, after categorization training, the top-down input from the association area to the mixture area induced a cascade of changes in the network’s representations of the sample stimulus, which was clearly shown at both the single neuron and population levels.

### Sample-test similarity change in the comparison module

Having characterized changes in the network’s representations of the sample stimulus due to category learning, we considered the sample-test comparison process in our model (Fig. 6A) that is essential to the similarity judgement in CP studies (Goldstone, 1994). In principle, comparison can be realized in different ways (Carpenter and Grossberg, 1987). For example, comparison can be realized by a single neural population that performs a simple addition of the sample and test inputs (Carpenter and Grossberg, 1987). Then, a match or non-match decision is determined by whether the sum exceeds a threshold. Recent work pointed out that this model fails with varying input magnitudes (Engel and Wang, 2011). Inspired by physiological observations (Miller et al., 1996; Freedman et al., 2003), a network architecture with two (ME and MS) neural populations (Fig. 6A) provides a robust and flexible way to make the sample-test comparison (Engel and Wang, 2011).

**Figure 6:**
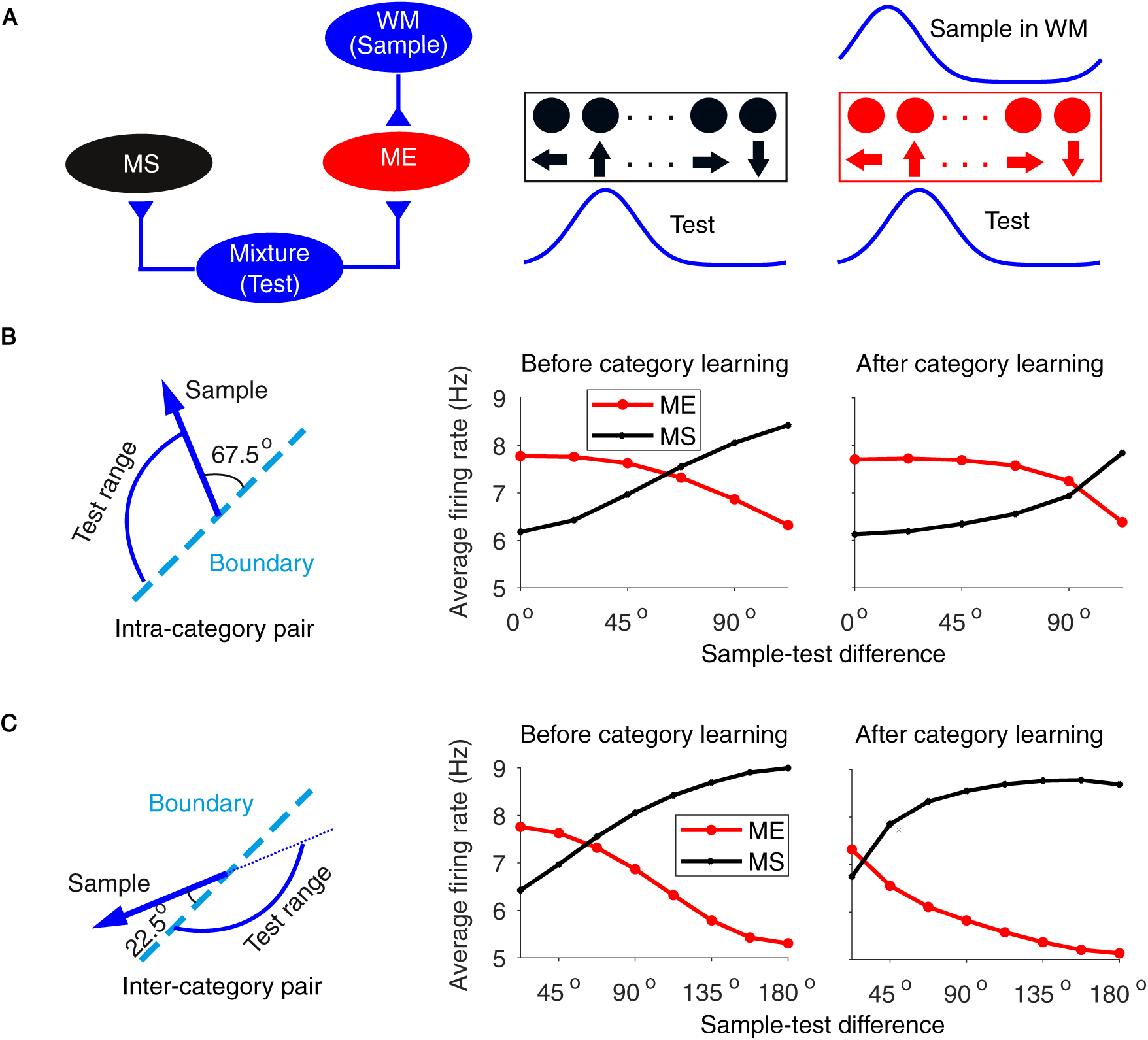
A similarity-based pattern match mechanism that can detect the similarity change induced by category learning. **(A)** Schematic of similarity-based pattern match mechanism in the comparison module. This module consists of two mutually inhibited functional populations – match enhancement (ME, red population) and match suppression (MS, black population), in which the ME population receives both top-down input from the working memory (WM) module and bottom-up input from the mixture area while the MS population only receives bottom-up input from the mixture area. (**B**) Average population firing rate of both ME (red line) and MS (black line) populations during the test period for intra-category pairs. The left panel depicts the configuration of intra-category pairs. (**C**) Average population firing rate of both ME (red line) and MS (black line) populations during the test period for inter-category pairs. The left panel depicts the configuration of inter-category pairs.

Generally, in this two-population architecture, higher ME population activity than MS population activity favors the match decision, and the opposite holds for the non-match decision. Therefore, it is important to quantify how neural firing rates of both ME and MS populations parametrically vary with the similarity between the sample and test activation patterns, which in our model is determined by a combination of inhibition-dominated recurrent dynamics in the comparison module and heterogeneous top-down modulation from the WM module (Engel and Wang, 2011). Specifically, the average neural firing rate of ME population should decrease as the sample-test directional difference increases, while the average neural firing rate of MS population should show the opposite trend (Fig. 6B). This scaling of comparison effects with sample-test directional difference has been found in prefrontal neurons in a non-human primate experiment (Hussar and Pasternak, 2012), supporting this similarity-based pattern match mechanism for the comparison process.

While this similarity-based comparison circuit worked well in the DMS task before category learning (Engel and Wang, 2011), it is unclear if the same comparison circuit can effectively capture the similarity change induced by categorization training. To address this question, we considered the similarity representation of intra-category pairs (Fig. 6B). As expected, before categorization training, the average firing rate of the ME population decreased as the sample-test directional difference increased, while the average firing rate of the MS population showed an opposite trend. After categorization training, the average firing rate of the ME population still decreased as the sample-test difference increased, but much less abruptly, while the average firing rate of the MS population still increased as the sample-test difference increased, but also much less abruptly. Therefore, given the same intra-category sample-test pair, compared to the case before categorization training, the average firing rate difference between ME and MS populations after categorization training became higher. This implies enhanced similarity for intra-category pairs after categorization training. For inter-category pairs, we found that, compared with the case before category learning, as the sample-test directional difference increased, the dependence of the average firing rate on the sample-test difference became steeper for both the ME and MS populations (Fig. 6C). Therefore, given the same inter-category sample-test pair, compared to the case before categorization training, the average firing rate difference between ME and MS populations after categorization training became smaller. This implied enhanced dissimilarity for inter-category pairs after categorization training. Therefore, this similarity-based comparison circuit in our model was capable of capturing both the intra- and inter-category similarity changes induced by categorization training.

### Match between model performance and monkey behavioral data

In our model, the analog sample-test similarity information was further transformed into a binary choice (match or non-match) in the decision module. This was achieved by attractor dynamics that amplified the difference between ME and MS inputs through slow recurrent excitation and feedback inhibition within the decision module (Wang, 2002). It has been shown that the stochastic choice behavior generated by attractor dynamics can be well-characterized by a sigmoid function of the difference in input currents (Soltani and Wang, 2006). More specifically, the fraction of choosing non-match over multiple trials can be approximated by a sigmoid function of the difference in input currents to non-match and match selective neural pools in the decision module (Fig. 7A). In the model, projection patterns from the comparison module to the decision module were learned through experience. Here, as in a previous work (Engel and Wang, 2011), reward-modulated Hebbian plasticity (for details, see Materials and Methods) was applied to the synaptic connections from the comparison module to the decision module. This gave rise to a connectivity pattern in which ME and MS neurons preferentially targeted match and non-match neuron pools, respectively (Fig. 7A).

**Figure 7:**
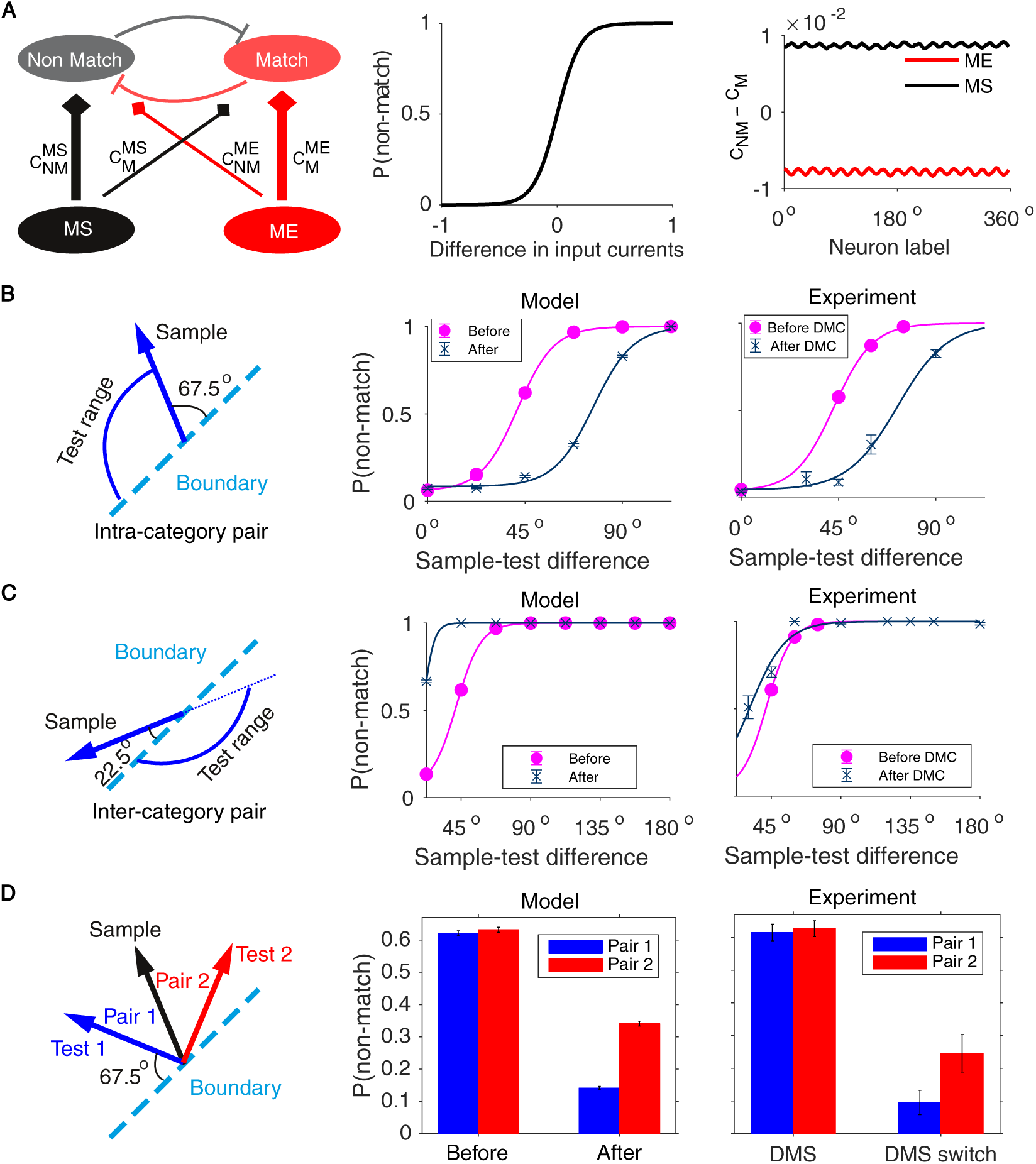
The circuit model is capable of accounting for the CP effect observed in the experiment. **(A)** Schematic of connections from the comparison module (with the ME and MS populations) to the decision module (with the non-match and match populations) (left panel). The stochastic choice behavior of the decision module can be characterized by a sigmoid function of the difference in input currents (middle panel). Right panel shows the difference between the ME-to-nonmatch synaptic weight and the ME-to-match synaptic weight (red curve) and the difference between the MS-to-nonmatch synaptic weight and the MS-to-match synaptic weight (black curve) as a function of neuron label. It can be observed that the ME population tends to connect with the match population more strongly while the MS population tends to connect with the non-match population more strongly. (**B-C**) The comparison of the DMS task performance for intra-category (B) and inter-category (C) pairs between model and experiment. The left panel depicts the configuration of sample and test directions. With the configuration in the left panel, the corresponding right panel shows how the fraction of reported non-match changes as a function of sample-test directional difference. (**D**) The comparison of two intra-category pairs between model and experiment. Error bars indicate s.e.m.

With this connectivity pattern, as the average firing rate of the MS (ME) population increased, the input currents to non-match (match) pools increased. As presented above, for both intra- and inter-category pairs before and after categorization training, as the sample-test directional difference increased, the average firing rate of the MS (ME) populations increased (decreased) (Fig. 6B and C). As a result, the difference in current inputs to non-match and match pools monotonically increased with respect to the sample-test directional difference. Since the choice behavior is a sigmoid function of the difference in input currents, the fraction of choosing non-match increased as sample-test directional difference increased in our simulations (Fig. 7B and C).

To gain insights into the question of how categorization training affects discrimination performance, we compared discrimination performance before and after categorization training. For intra-category pairs, compared to the case before categorization training, the fraction of choosing non-match after categorization training in our model was significantly lower, recapitulating the compression effect in the experimental data (Fig. 7B). To explain this result in the model, we considered an intra-category sample-test pair with sample and test stimuli 45° apart. For this pair, compared to the case before categorization training, the average firing rate of the MS (ME) populations became lower (higher) (Fig. 6B). Given the fixed synaptic weights from the comparison module to the decision module, this meant a smaller difference in current inputs to the non-match and match pools after categorization training, leading to a smaller fraction of choosing non-match after categorization training. For inter-category pairs, compared to the case before categorization training, the fraction of choosing non-match after categorization training in our model was significantly higher, recapitulating the expansion effect in the experiment (Fig. 7C). Similarly, let us consider an inter-category sample-test pair with sample and test stimuli 45° apart. For this pair, compared to the case before categorization training, the average firing rate of the MS (ME) populations became higher (lower) (Fig. 6C). This meant a bigger difference in current inputs to the non-match and match pools after categorization training, leading to a larger fraction of choosing non-match after categorization training. Furthermore, our model can also recapitulate the discrimination performance difference for pairs with the same sample-test directional difference but different distances from the category center found in the experiment (Fig. 7D). In summary, our model was capable of accounting for the behavioral data in the experiment above and provided a potential mechanistic explanation for it.

### The cognitive benefit of having CP

Given the ubiquity of CP effects in different domains, different hypotheses regarding the cognitive function of having CP have been proposed (Goldstone and Hendrickson, 2010). Here, we tested the hypothesis that category knowledge acquired during category learning can help improve perceptual stability in noisy environments. To this end, we varied the sensory input amplitude to model the environmental change. The smaller the input amplitude is, the noisier the environment is. Meanwhile, we used the average firing rate of neurons in the mixture area to characterize perceptual stability. The higher the average firing rate is, the more stable the perception is. Furthermore, the average CTI value of neurons in the mixture area was computed to quantify how strong the mixture area is tuned to categories. The larger the CTI is, the stronger the mixture area is tuned to categories.

From the middle panel of Fig. 8A, we can observe that the average firing rate of neurons in the mixture area after category learning is higher than that before category learning, indicating improved perceptual stability after category learning. Interestingly, the right panel of Fig. 8A shows that the average CTI value of neurons in the mixture area increases as the input amplitude decreases. This suggests that category knowledge plays an increasingly important role in stabilizing sensory perception as the environment become noisier. Since the category information is purely sent by the feedback projections from the association area to the mixture area, this result indicates the importance of feedback projections in improving perceptual stability in noisy environments. To address the functional importance of feedforward projections from the sensory area to the mixture area, we removed the feedforward projections and examined the resulting firing patterns of neurons in the mixture area. We found that in this scenario while category learning can still improve perceptual stability (the middle panel of Fig. 8B), the CTI value becomes extremely high all over the tested input amplitude range (the right panel of Fig. 8B), which could be detrimental to precise feature-based sensory perception. Therefore, while feedback category input can improve perceptual stability in environments with high uncertainty (i.e., small sensory input amplitude), the feedforward sensory input is also required to enable precise featured-based perception in environments with low uncertainty (i.e., large sensory input amplitude). In this sense, the existence of a mixture area can be regarded as a natural way to implement the balance between keeping perceptual stability in environments with high uncertainty and maintaining perceptual sensitivity in environments with low uncertainty. Therefore, the CP effect can be understood as an emerging phenomenon resulting from this intriguing balance.

**Figure 8:**
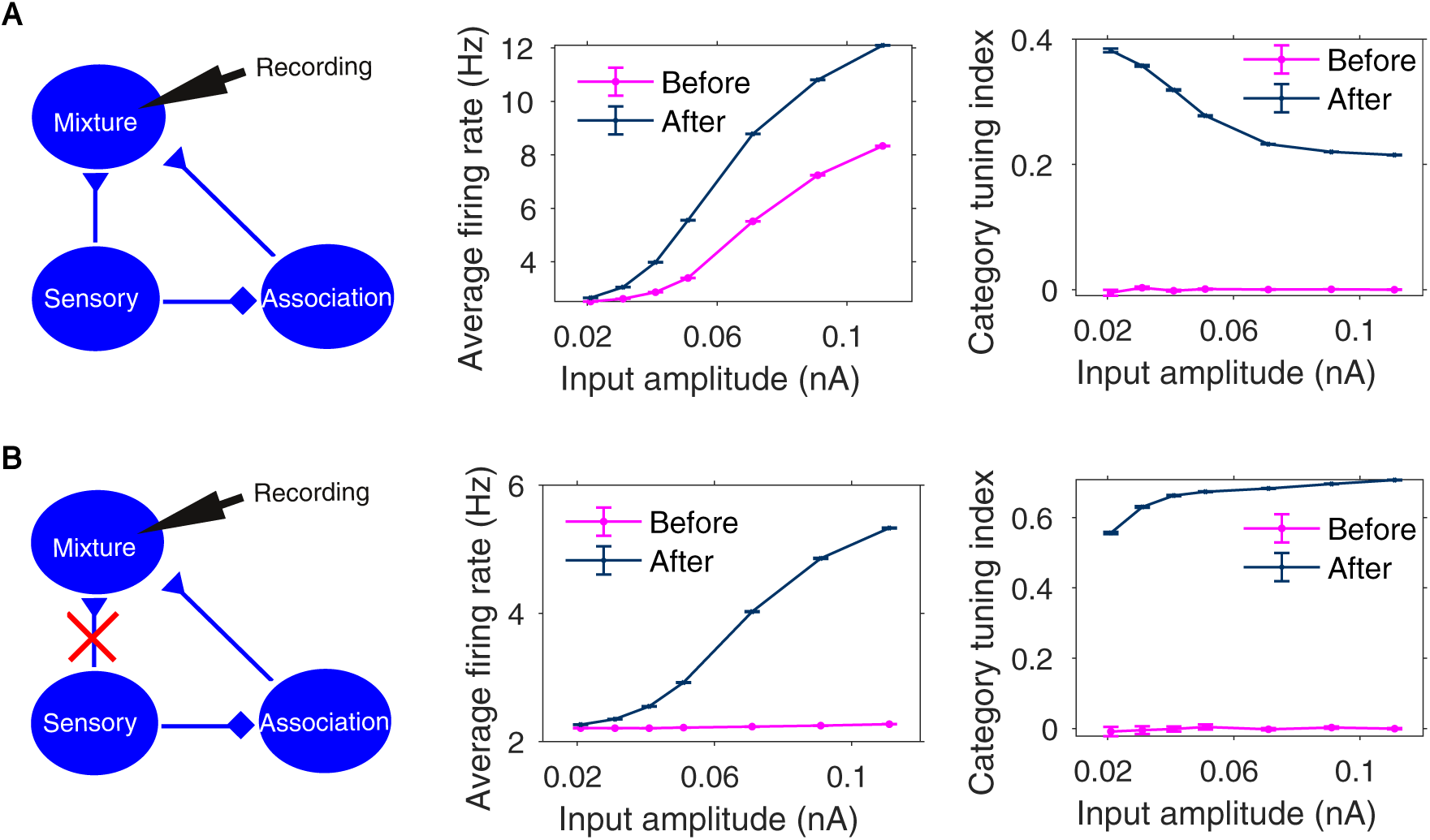
The cognitive benefit of having CP – improving perceptual stability in noisy environments with salient category information. **(A)** Left panel: schematic of connections within the sensory module of our original model. Middle panel: given the sample direction at the category center, the average firing rate of neurons in the mixture area as a function of input amplitude during the sample period before and after category learning. The average firing rate after category learning is higher than that before category learning, indicating improved perceptual stability after category learning. Right panel: the average category tuning index (CTI) of neurons in the mixture area as a function of input amplitude during the sample period before and after category learning. The average CTI value becomes non-zero after category learning. In particular, the CTI value after category learning increases as input amplitude decreases, suggesting that the feedback category input plays an increasingly important role in stabilizing sensory perception for small input amplitudes. (**B**) Scenario in which feedforward projections from the sensory area to the mixture area are removed from our model. In this scenario, both the average firing rate and the CTI value after category learning become higher than the corresponding ones before category learning. However, the CTI value becomes very high, which could be detrimental to precise feature-based sensory perception. Therefore, feedforward projections from the sensory area to the mixture area are also functionally important. Error bars indicate s.e.m. for both (**A**) and (**B**).

## Discussion

CP is a key cognitive process revealing the intriguing interplay between analog feature-based perception and discrete categorization. However, its neural mechanisms have been largely unknown. Here, we showed behavioral evidence for the learned CP effect in a visual motion delayed matching experiment in non-human primates. By leveraging existing key neurophysiological findings in visual categorization, working memory and decision making, we developed a computational framework that allowed us to make testable predictions for further experiments. For example, our model predicted the existence of a mixture area that integrates feedforward sensory input and feedback category input. Furthermore, by showing that category learning enables improved perceptual stability in noisy environments, our work provided a functional explanation for the existence of CP phenomenon. Taken together, our work suggests that visual motion delayed matching paradigms in non-human primates combined with biologically-based modeling can serve as a promising model system for elucidating the neural mechanisms of learned CP phenomenon.

As an important step towards elucidating the neural mechanisms of CP effect, the neural basis of categorization has drawn much attention recently. Studies of visual categorization in nonhuman primates suggest that changes in neural representations due to category learning occur in higher-order visual areas (Sigala and Logothetis, 2002; Sigala et al., 2002) and higher association cortex (Freedman and Assad, 2006; Swaminathan et al., 2013; Sarma et al., 2016), but not earlier sensory cortex (Freedman and Assad, 2006). This line of research provides important insights into the question of how the category knowledge acquired during categorization is stored in our brain. However, recent studies in humans and rodents indicate that category biases during categorization tasks can occur in signals in earlier visual and auditory cortices (Ester et al., 2020; Xin et al., 2019). Whether this reflects a species difference, signal measurement difference (Mendoza-Halliday et al., 2014) (spiking activity, functional MRI or EEG) or the importance of the training protocol (Ester et al., 2020) (for example, human subjects learned to categorize stimuli after about 10 minutes of training, as opposed to over periods of several months in the monkey studies) remains to be established.

In the realm of learned CP, efforts have been made to quantify the degree to which category learning influences perceptual discrimination in human psychophysical studies, with conflicting results (Goldstone, 1994; Livingston et al., 1998; Jiang et al., 2007; Gureckis and Goldstone, 2008; Folstein et al., 2012). Some studies showed that objects became more discriminable along dimensions relevant to the learned categories (Goldstone, 1994; Gureckis and Goldstone, 2008), while others did not find this effect (Jiang et al., 2007). In this work, we showed a distance-dependent CP effect along the dimension relevant to the learned categories in a delayed-matching-type discrimination task in non-human primates. It would be of interest to test the robustness of this CP effect by using different perceptual discrimination tasks.

Given the ubiquity of behavioral evidence for CP, various computational models of CP have been developed, including Bayesian models (Kronrod et al., 2016; Feldman et al., 2009) and connectionist neural network models (Damper and Harnad, 2000). By contrast, our biophysical model was grounded in key neurophysiological findings (Freedman and Assad, 2016; Leavitt et al., 2017; Hussar and Pasternak, 2012; Gold and Shadlen, 2007) and biological learning rules (Schultz et al., 1997; Schultz, 2007; Fremaux et al., 2010; Loewenstein and Seung, 2006; Fusi et al., 2007; Soltani and Wang, 2010). This enabled us to make experimentally testable predictions regarding neurophysiological evidence and anatomical substrate for CP. For example, in this work, we hypothesized that learned category knowledge gets into the perceptual system as early as during the sensory encoding stage through feedback projectionsfrom the association area to the mixture area. To experimentally test this hypothesis, it would be important to identify the anatomical correspondence of the mixture area. While it seems challenging, several clues allude to the possibility that MST area could be one candidate for the mixture area. First, during a visual categorization task, neuronal categorical representations emerged in the LIP area, but not in the MT area, during the sample period (Freedman and Assad, 2006). This suggested that if there is any area showing neuronal categorical representations during the sample period, the area must be higher than MT in the cortical hierarchy. As an area immediately downstream to MT that processes visual motion information, MST is well-positioned to be this area. Second, during a visual motion working memory task, neurons in MST showed strong sustained spiking activity encoding memorized visual motion directions, in sharp contrast to neurons in MT showing no sustained spiking activity (Mendoza-Halliday et al., 2014). This indicated that MST may be markedly different from MT in serving high cognitive function, including working memory and categorization. Therefore, it would be of interest to look for the physiological signatures of neuronal categorical representations (Fig. 5D) in MST during the DMS task after category learning.

In our model, the sensory encoding change eventually would manifest itself during all stages in the downstream information processing. Among these stages, the comparison stage is of particular interest because the anatomical correspondence of the comparison module has been relatively well-studied in the literature. For example, in a memory-guided motion direction discrimination task (Hussar and Pasternak, 2012), neural activity in prefrontal cortex exhibited substantial comparison effects that were consistent with the proposed similarity-based pattern match mechanism (Engel and Wang, 2011). In analyses of experimental data, the area under the receiver operating characteristic curve (AROC) – a metric within signal detection theory – was used to quantify the same-different comparison effect of both ME and MS neurons (Hussar and Pasternak, 2012). The smaller the AROC is, the more similar the sample and test stimuli are. For stimulus pairs with the same sample-test directional difference, our model predicted that the intra-category pair will produce more similar ME and MS neuron activity than the inter-category pair. ME and MS neurons are thought to exist in the prefrontal cortex. Hence, in PFC, the measured AROC of the intra-category pair should be smaller than that of the inter-category pair during the DMS task after category learning, which is experimentally testable.

In this work, we demonstrated the importance of feedback signals from a category circuit (the association area) in improving perceptual stability in noisy environments. Interestingly, a previous modeling work showed that the similar category circuit plays a key role in resolving sensory-motor conflict in a rule-based switching task (Ardid and Wang, 2013). Taken together, this line of modeling work provides concrete hypotheses about how category learning can facilitate our cognitive performance (Seger and Miller, 2010).

It has been elusive to dissect physiological fingerprints and functional roles of feedback projections in the cortical hierarchy. To summarize, the present work combining experiments and modeling opens the door for the elucidation of top-down signaling in the cortex of an especially important kind, namely influence of sensory processing by abstract (category) knowledge.

## Acknowledgements

The authors thank N. Santhanam, R. Kharkar and M. Rose for assistance with animal training, J. Hodnefield for expert behavioral and experimental assistance, S. Thomas for expert technical assistance, C. Gray for technical and surgical assistance, and the staff of The University of Chicago Animal Resources Center for expert veterinary assistance. We also thank W. Pettine for valuable comments on an earlier version of this manuscript. This work was supported by grants from the National Science Foundation (NSF 1631586 to X.-J.W. and D.J.F.). Additional support was provided by a McKnight Scholar award, the Alfred P. Sloan Foundation, NSF CAREER award 0955640, the Brain Research Foundation, NIH R01EY019041 (D.J.F.), NRSA MH097428 (A.S.), R01-MH062349, 5U19NS107609 (X.-J.W.) and NSFC 11901557 (B.M.).

## Author Contributions

B.M., D.P.B., D.J.F., and X.-J.W. designed the research. B.M. performed the data analysis, modeling work and wrote the manuscript. D.P.B. performed the data analysis and edited the manuscript. A.S. trained the animals and performed the data analysis. D.J.F. supervised the experimental work, assisted with animal training, data analysis and edited the manuscript. X.-J.W. supervised the modeling work and data analysis, and edited the manuscript.

